## Supplementary data for "*Drosophila* larval models of invasive tumourigenesis for in vivo studies on tumour/peripheral host tissue interactions during cancer cachexia"

1. CRUK Beatson Institute. Institute of Cancer Sciences. University of Glasgow. Garscube Estate, Switchback Road, G61 1BD. Glasgow, U.K.
2. Wolfson Wohl Cancer Research Centre. Institute of Cancer Sciences. University of Glasgow. Garscube Estate, Switchback Road, G61 1QH. Glasgow, U.K.

‡ Equally contributing authors

#### Supplementary Figure Legends

**Figure S1. Representative images of imaginal disc tissues. (A,C,E-L)** Immunofluorescent confocal images of larval imaginal disc stained with DAPI (blue) from control (*$w^{1118}$*) **(A)** animals carrying hyperplastic (*rotund>$Yki^{S168A}$*) tumours **(C)**, non-invasive neoplastic tumour (*$dlg^{40.2}$*) **(E, F),** malignant neoplastic MARCM-generated eye tumours (*$Ras^{V12}$*, *$scrib^{1}$*) **(G, H)** or *hedgehog-gal4* overexpressing *$Ras^{V12}$*, *$scrib^{IR}$* (*$Ras^{V12}$*, *$scrib^{IR}$*) tumours at 6 or 12 days AED. Imaginal discs from 6- or 12-days AED *$e^{1}$* larvae **(K, L). (**B) Brightfield image of *$w^{1118}$* adult fly at 12 days AED. (D) Brightfield image of dead *rotund>$Yki^{S168A}$* pupa at twelve days AED.

**Figure S2. Representative images of animals and larval cuticles. (A,C,E-L)** Immunofluorescent confocal images of body wall muscle from animals of genotypes and ages as indicated in Figure S1. **(B)** Brightfield image of *$w^{1118}$* adult fly at 12 days AED. **(D)** Brightfield image of dead *$Yki^{S168A}$* pupa at twelve days AED.

**Figure S3. Representative images of the 5$^{th}$ segment of the 7$^{th}$ ventral muscle. (A-F,I-L)** Immunofluorescent confocal images of the 5$^{th}$ segment of the 7$^{th}$ ventral larval muscle (outlined in yellow in the grey scale images) from animals of genotypes an ages as indicated in Figure S1. Tissues were stained with Phalloidin (green) and LipidTOX (red and grey). **(G)** Brightfield image of *$w^{1118}$* adult fly at 12 days AED. **(H)** Brightfield image of dead *$Yki^{S168A}$* pupa at 12 days AED. The panels denoted with (‘) represent lipid staining alone. The panels denoted with (“) represent a transversal orthographic view of the muscle shown in (A-F,I-L), with a 2X zoom. Areas demarked by red boxes represent zoomed in images in insets.

**Figure S4. Impl2 does not mediate muscle wasting in larval models of tumour induced cachexia. (A)** mRNA expression levels of *impl2* in skeletal muscle from 7-day-old wild type larvae (*$w^{1118}$*) or *hedgehog-gal4* overexpressing *$Ras^{V12}$*, *$scrib^{IR}$* tumours without (*$w^{1118}$*) or with *impl2* knockdown (*impl2-IR).* One-way ANOVA with Tukey post-hoc correction. *p≤0.05; ***p≤0.001. (B) Quantification of the percentage of cuticle covered by muscle in 12-day-old larvae with *hedgehog-gal4* overexpressing $Ras^{V12}$, $scrib^{IR}$ tumours without ($w^{1118}$) or with *impl2* knockdown (*impl2-IR)*. T-test; no statistical significance.

**Supplementary Table Legends**

**Table S1: RNAseq data of cuticles from control and cachectic animals.** Complete data set from RNAseq analysis of cuticles from control and cachectic animals (Figure 4). Control condition is labelled in blue and named 'cont', while the data corresponding to cachectic animals is labelled in red and named 'treat'.

**Table S2: Fly lines and full genotypes of animals used throughout the study.**

**Table S3: Sequences of primer sets used throughout the study.**

### Figure S1

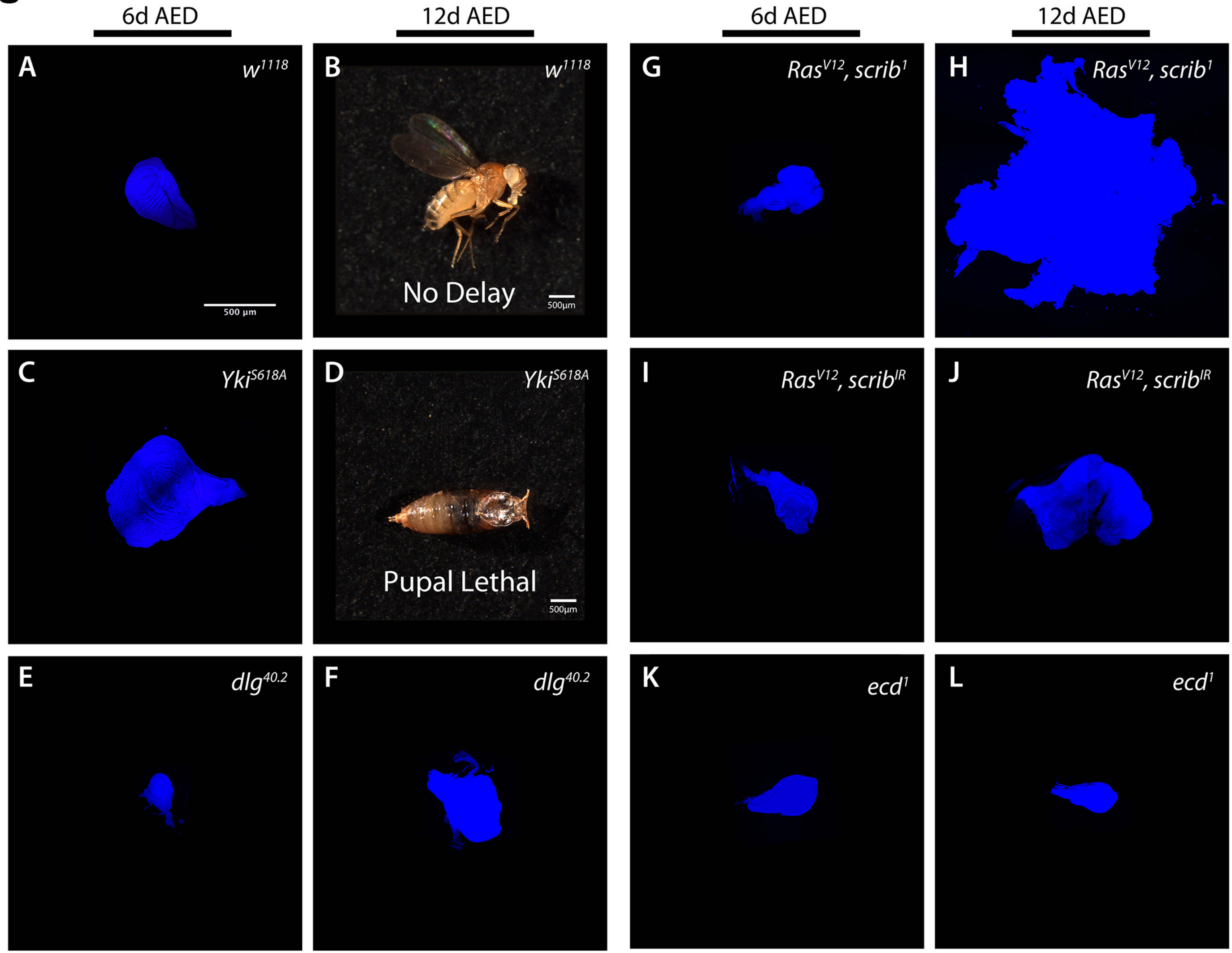
6d AED
12d AED
6d AED
12d AED
A
w1118
500 µm
B
w1118
No Delay
500µm
G
RasV12, scrib1
H
RasV12, scrib1
C
YkiS618A
D
YkiS618A
Pupal Lethal
500µm
I
RasV12, scribIR
J
RasV12, scribIR
E
dlg40.2
F
dlg40.2
K
ecd1
L
ecd1

### Figure S2

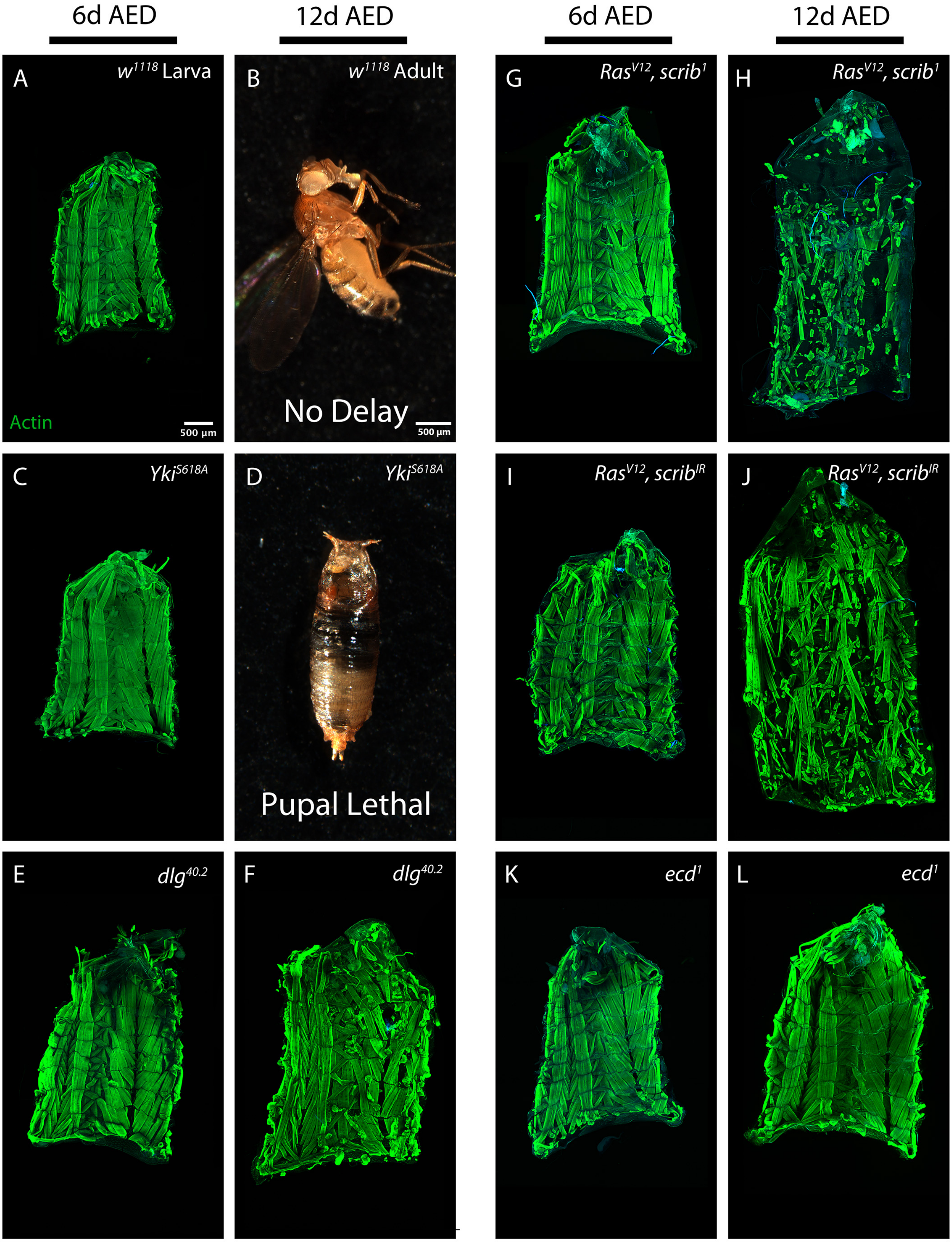

6d AED
12d AED
6d AED
12d AED
A
w1118 Larva
Actin
500 µm
B
w1118 Adult
No Delay
500 µm
G
RasV12, scrib1
H
RasV12, scrib1
C
YkiS618A
D
YkiS618A
Pupal Lethal
I
RasV12, scribIR
J
RasV12, scribIR
E
dlg40.2
F
dlg40.2
K
ecd1
L
ecd1

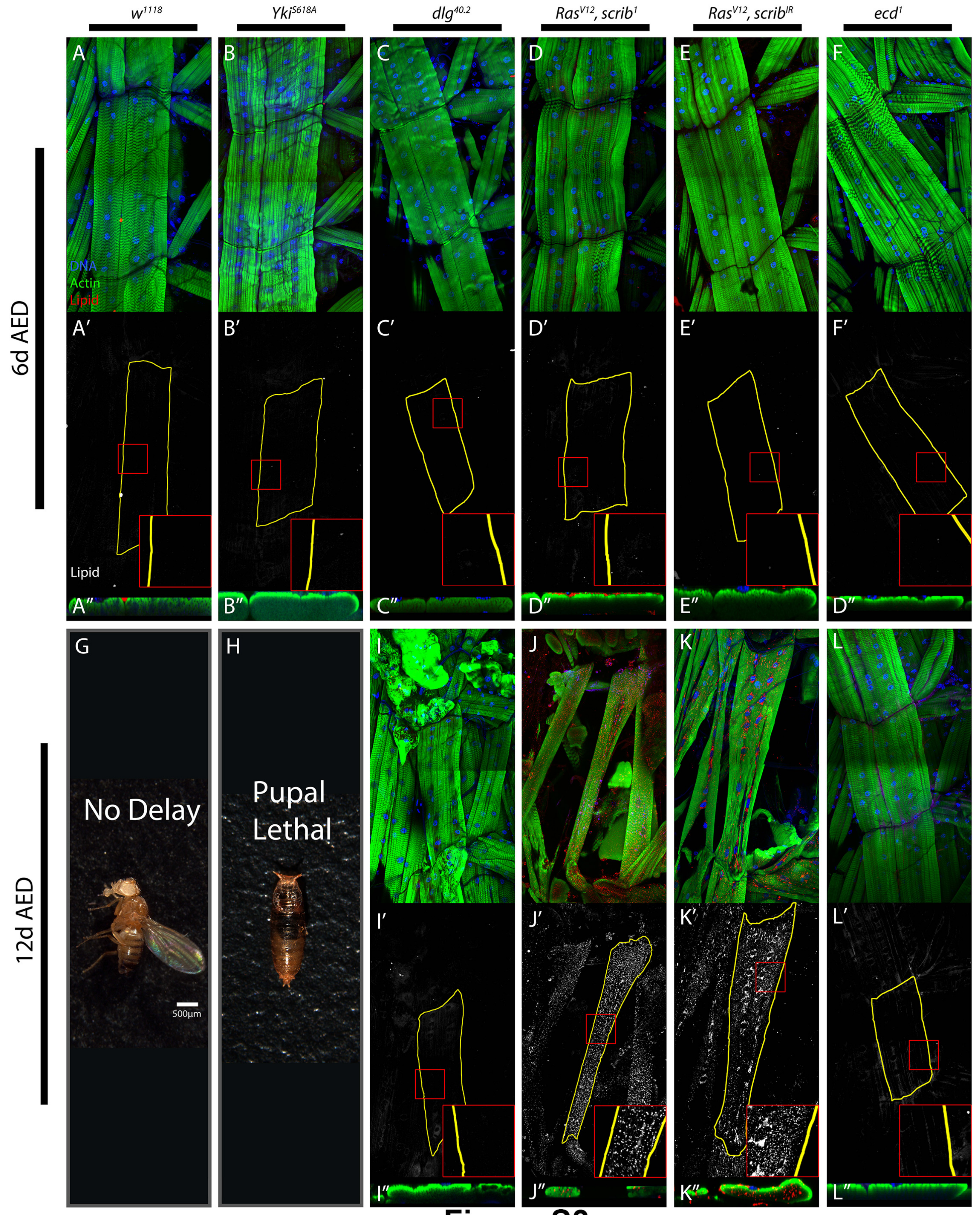

w1118
YkiS618A
dlg40.2
RasV12, scrib1
RasV12, scribIR
ecd1
6d AED
12d AED
A
B
C
D
E
F
DNA
Actin
Lipid
A′
B′
C′
D′
E′
F′
Lipid
A″
B″
C″
D″
E″
D″
G
H
I
J
K
L
No Delay
Pupal Lethal
500µm
I′
J′
K′
L′
I″
J″
K″
L″


### Figure S3

### Figure S4

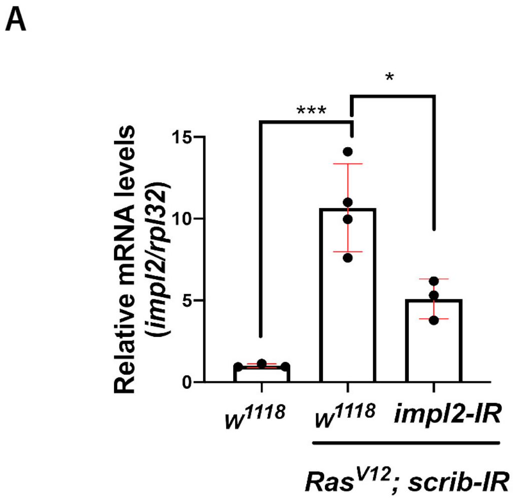
A
15
10
5
0
Relative mRNA levels
(impl2/rpl32)
***
*
w1118
w1118
impl2-IR
RasV12; scrib-IR

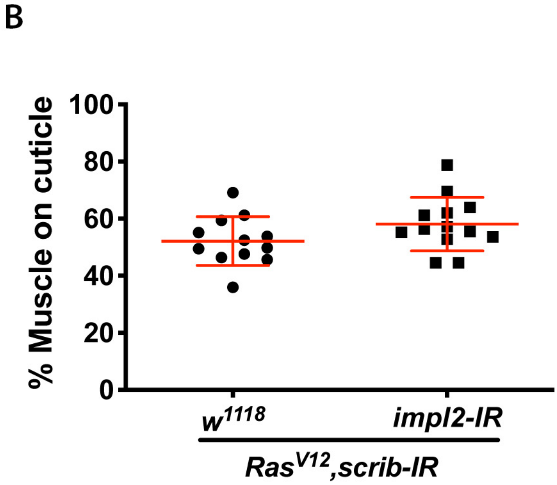
B
100
80
60
40
20
0
% Muscle on cuticle
w1118
impl2-IR
RasV12,scrib-IR
